# Different genetic determinants for high virulence, transmission and replication of high pathogenicity H7N7 avian influenza virus in turkeys and chickens

**DOI:** 10.1101/2025.03.18.643940

**Authors:** David Scheibner, Juliane Lang, Christine Luttermann, Diana I. Palme, Kati Franzke, Dajana Helke, Maryna Kuryshko, Olanrewaju I. Fatola, Mathilde Richard, Jutta Veits, Thomas C. Mettenleiter, Reiner Ulrich, Elsayed M. Abdelwhab

## Abstract

High pathogenicity (HP) avian influenza viruses (AIV) generally evolve from low pathogenicity (LP) precursors after transmission from wild birds to chickens (*Gallus gallus domesticus*) and turkeys (*Meleagris gallopavo*), causing severe economic losses worldwide. Turkeys are more susceptible to AIV infection than chickens and are considered potential bridging hosts that facilitate the emergence of HPAIV. Beyond the polybasic cleavage site (pCS) in hemagglutinin (HA), little is known about other virulence determinants of HPAIV in these species. In 2015, HPAIV H7N7 and its LP ancestor were isolated from the same chicken farm, which differed by 16 nonsynonymous mutations across all eight gene segments, in addition to the pCS. Here we identify the genetic determinants, including the pCS, that contributed to the HPAIV H7N7 virulence, transmission, replication, and tissue distribution in chickens and turkeys. Notably, the non-structural (NS1) or matrix (M) proteins’ encoding segments in turkeys, or NS segment in chickens, rendered viruses as virulent and transmissible as the original HPAIV. Endotheliotropism, observed exclusively in chickens, was driven by the pCS and, to a lesser extent, the neuraminidase (NA). In vitro, the M2-V68L mutation influenced NS1 expression and virus morphology in chicken and turkey cells. Additionally, HPAIV NS1 enhanced polymerase activity and effectively suppressed interferon induction, a process further modulated by M2-V68L. These findings underscore the critical role of turkeys as a “hub” in the evolution of HPAIV from LP precursors, offering crucial insights into the genotypic and phenotypic factors that facilitate viral adaptation in different poultry species.

**Importance:** High pathogenicity avian influenza viruses (HPAIV) cause severe economic losses for the poultry industry worldwide. HPAIV generally evolve from low pathogenicity (LP) ancestors in galliform birds, with turkeys being more susceptible to severe disease and death than chickens. The mechanisms underlying HPAIV emergence in these species remain unclear. This study reveals two distinct evolutionary pathways for HPAIV. In turkeys, both the polybasic hemagglutinin cleavage site (pCS) and mutations in the NS or M segments contributed to high virulence and transmission. In chickens, only the NS segment was critical, in addition to the pCS. These segments increased virus replication in both chicken and turkey cells. However, unlike chicken cells, the M and NS segments did not play a role in blocking the innate immune response. Understanding these species-specific mechanisms highlights the role of turkeys as a bridging host and provides insights into the molecular evolution of HPAIV from LP precursors.

## Introduction

Avian influenza viruses (AIV) of the genus Influenza A Virus (IAV) in the family *Orthomyxoviridae* possess a single-stranded RNA genome composed of eight gene segments encoding at least ten structural and 1 non-structural proteins. These include polymerase basic subunit 2 (PB2) protein, polymerase basic subunit 1 (PB1) protein, polymerase acidic (PA) protein, hemagglutinin (HA), nucleoprotein (NP), neuraminidase (NA), matrix proteins M1/M2, nuclear export protein (NEP) and the non-structural (NS) protein NS1 (1). The genome of some AIV encode also additional non-structural proteins like PB1-F2, and PA-x (2). These proteins are essential for the viral replication cycle. For example, HA mediates cell entry via binding the sialic acid receptors, and should be cleaved into HA1 and HA2 subunits by host proteases to facilitate fusion, a process that is pH-dependent and regulated by the M2 ion channel. NS1 antagonizes interferon responses, while the ribonucleoprotein complex (PB2, PB1, PA, and NP) drives RNA replication and transcription. M1 facilitates virus assembly, and NA cleaves sialic acids to enable progeny virion release (1).

AIV subtypes are classified by their HA and NA antigenic properties, with 17 HA and 9 NA subtypes found in wild bird reservoirs. While all H1-H16/H19 subtypes can infect birds, only H5 and H7 subtypes may evolve into high pathogenicity (HP) forms, causing up to 100% mortality in poultry (3–6). Pathogenicity of H5 and H7 viruses is largely determined by the amino acid (aa) sequence of the HA proteolytic cleavage site (CS), which influences tissue distribution and virulence. Low pathogenicity AIVs (LPAIV) possess a monobasic CS cleaved by trypsin-like proteases, restricting virus replication in respiratory and intestinal tracts. In contrast, HPAIV carry a polybasic CS (pCS), enabling activation by furin-like proteases abundant in various tissues, particularly in galliformes (e.g. chickens and turkeys)(6–10). HPAIV evolve from LPAIV through mutations in the HA CS and/or other gene segments with unique constellations influencing virulence and transmission in a strain-dependent manner (11–14). Therefore, it is important to study the virulence determinants of each HPAIV outbreak separately. Natural evolution of H7Nx HPAIV from LPAIV has been frequently documented in chickens and turkeys (rarely in ducks) (15). However, virulence determinants in poultry, particularly turkeys, remain poorly understood (11, 16).

Chickens and turkeys exhibit high mortality upon HPAIV infection. However, field and experimental findings indicate turkeys are more susceptible to AIV-induced morbidity and mortality than chickens (6, 16–19). The molecular mechanisms driving these differences, particularly for H7 HPAIV, remain inadequately investigated. Recently, we have demonstrated that the pCS alone was the main virulence determinant for H7N1 in turkeys, but not in chickens (20). In 2015, HPAIV H7N7 and its LPAIV precursor were isolated from the same laying-egg chicken farm in Germany (21). The LPAIV possessed a monobasic HACS ^315^PEIPKGR/GLF^324^, while the HPAIV had developed a polybasic motif ^315^PEIPKGR**KRR**/GLF^327^ and additional 16 nonsynonymous substitutions across multiple gene segments, including PB2 (E123K, I147V, K355R), PB1 (I246M, F254C), PA (K185R), HA (I13S in the signal peptide), NP (S478F), NA (P48K, V439A, I476T), M2 (V68L), NS1 (N92D, G209D), and NEP (V52M, K95E). We previously showed that the H7N1 LPAIV with only the pCS was less virulent in turkeys than in chickens, suggesting additional mutations contribute to virulence (22). Here, using reverse genetics, recombinant parental LPAIV but carrying the pCS (LP_poly) in combination with different gene segments of the subsequent HPAIV were generated to identify the minimal gene constellation for high virulence and transmission for both species. These viruses were tested in chickens and turkeys, with the mechanisms of variable virulence further analyzed *in vitro*.

## Results

### Recombinant H7N7 viruses

All gene segments of the LP and HP viruses as well as LP_poly were generated in a previous study (22). Additional recombinant viruses were created by combining LP_poly with individual HP-derived gene segments (designated LP_poly_[segment name]) or all three polymerase subunits (designated LP_poly_P3) (Table 1). These recombinant viruses achieved titres of 10^6^ to 10^7^ PFU/ml in allantoic fluids after propagation in embryonated chicken eggs (ECE) (Table 1). Sanger sequencing confirmed the absence of unwanted mutations at the consensus level. However, all attempts to generate a viable LP_poly_NP virus failed, excluding it from further experiments in this study.

**Table 1:** Recombinant viruses used in this study.

| No. | Abbreviation | Cleavage site | Segment from HP | Titer (log10 PFU/mL) | Used to inoculate |  |
| --- | --- | --- | --- | --- | --- | --- |
|  |  |  |  |  | Turkeys | Chickens |
| 1 | LP* | Monobasic | None | 7.0 | Yes | Yes |
| 2 | LP_poly* | Polybasic | None | 6.7 | Yes | Yes |
| 3 | LP_poly_PB2 | Polybasic | PB2 | 6.0 | Yes | No |
| 4 | LP_poly_PB1 | Polybasic | PB1 | 6.7 | Yes | No |
| 5 | LP_poly_PA | Polybasic | PA | 6.6 | Yes | Yes |
| 6 | LP_poly_P3 | Polybasic | PB2, PB1, PA | 6.0 | No | Yes |
| 7 | LP_HA | Polybasic | HA | 6.5 | Yes | Yes |
| 8 | LP_poly_NA | Polybasic | NA | 6.7 | Yes | Yes |
| 9 | LP_poly_M | Polybasic | M | 7.0 | Yes | Yes |
| 10 | LP_poly_NS | Polybasic | NS | 6.8 | Yes | Yes |
| 11 | HP* | Polybasic | All gene segments | 6.9 | Yes | Yes |

\* These viruses were generated in a previous study (Scheibner et al. 2021)

### In turkeys, M and NS segments, in addition to the pCS, were main determinants for high virulence

We inoculated turkeys with different viruses via the oculonasal route (ON) and added naïve turkeys to each group at 1-day post-inoculation (dpi) to assess direct turkey-to-turkey contact transmission. Birds were monitored daily to assess mortality rate, mean time to death (MTD), and pathogenicity index (PI), ranging from 0 (avirulent) to 3 (maximal virulence). All HP-inoculated turkeys died within six days, with an MTD of 5.4 days and a PI of 2.0. Co-housed turkeys died within 6 days post-contact (dpc) (Table 2, Figure 1A, B). LP_poly_M and LP_poly_NS were as virulent and transmissible as HP, with all inoculated turkeys dying with MTD 6.1 and 5.7 days and PI 1.8 and 2.0, respectively. All co-housed turkeys died within 7 dpc. Other LP_poly reassortants with HP PB2, PB1, PA, HA, or NA were less virulent, with PI values ranging from 0.6 to 1.3, and most birds survived (Table 2). Neurological signs, including depression, ataxia, and head tilt, were prominent in the LP_poly_M, LP_poly_NS, and HP groups. The amount of virus excretion from surviving birds were determined in oropharyngeal (OP) and cloacal (CL) swabs at 4 dpi and anti-NP antibodies were detected at the end of the experiment (10 dpi). Overall, inoculated and contact turkeys excreted comparable viral RNA amounts in the OP and CL swabs. Notably, LP_poly (and to a lesser extent HP) was detected at higher levels than LP in in the CL swabs of inoculated turkeys (p = 0.0369) (Figure 1C-F). All surviving inoculated and contact turkeys had seroconverted by the end of the experiment. These findings highlight that M and NS segments, in addition to the pCS, are key determinants for virulence of this H7N7 HPAIV in turkeys.

**Figure 1:**
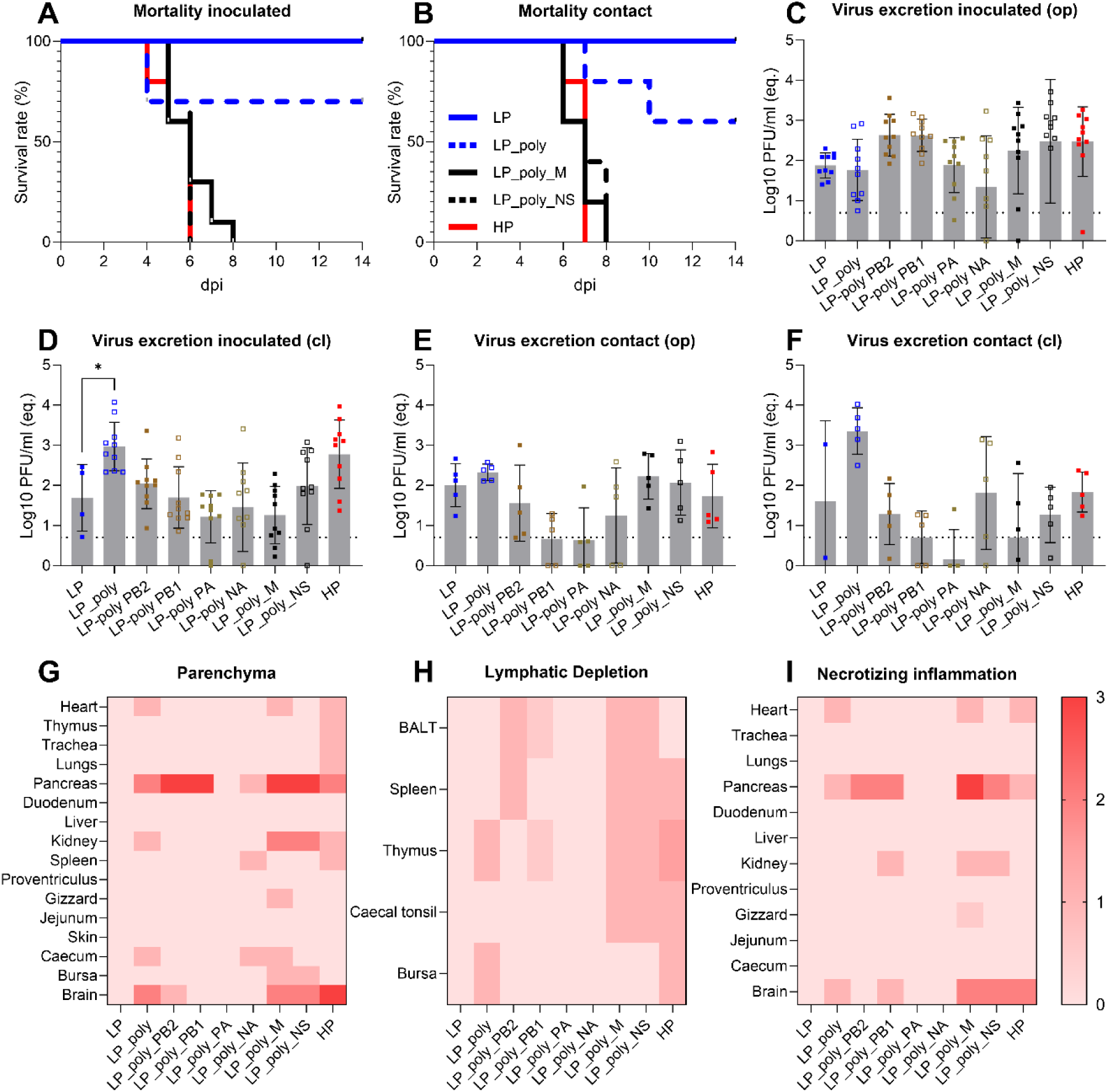
Clinical examination, virus excretion, tissue distribution and microscopic lesion in turkeys. Shown are the mortality of inoculated (A) and contact turkeys (B), virus RNA levels expressed as equivalent log10 plaque forming units/ml in oropharyngeal (OP) and cloacal (CL) swabs at 4 dpi from inoculated (C,D) and contact turkeys (E,F). Bars indicate the arithmetic mean of log-transformed data, with individual values represented as dots and standard deviations shown as error bars. The RT-qPCR detection limit is marked by dashed lines. Statistical differences in virus excretion compared to LP-inoculated birds were calculated with Kruskal-Wallis test and p<0.05 was considered significant (C-F). The median score for distribution of M1 antigen in inoculated turkeys at 4 dpi in the parenchyma using immunohistochemistry (G), necrosis in lymphatic organs and tissues (H) and severity of necrotizing inflammation (I). No viral antigens were detected in the endothelial cells.

**Table 2:**
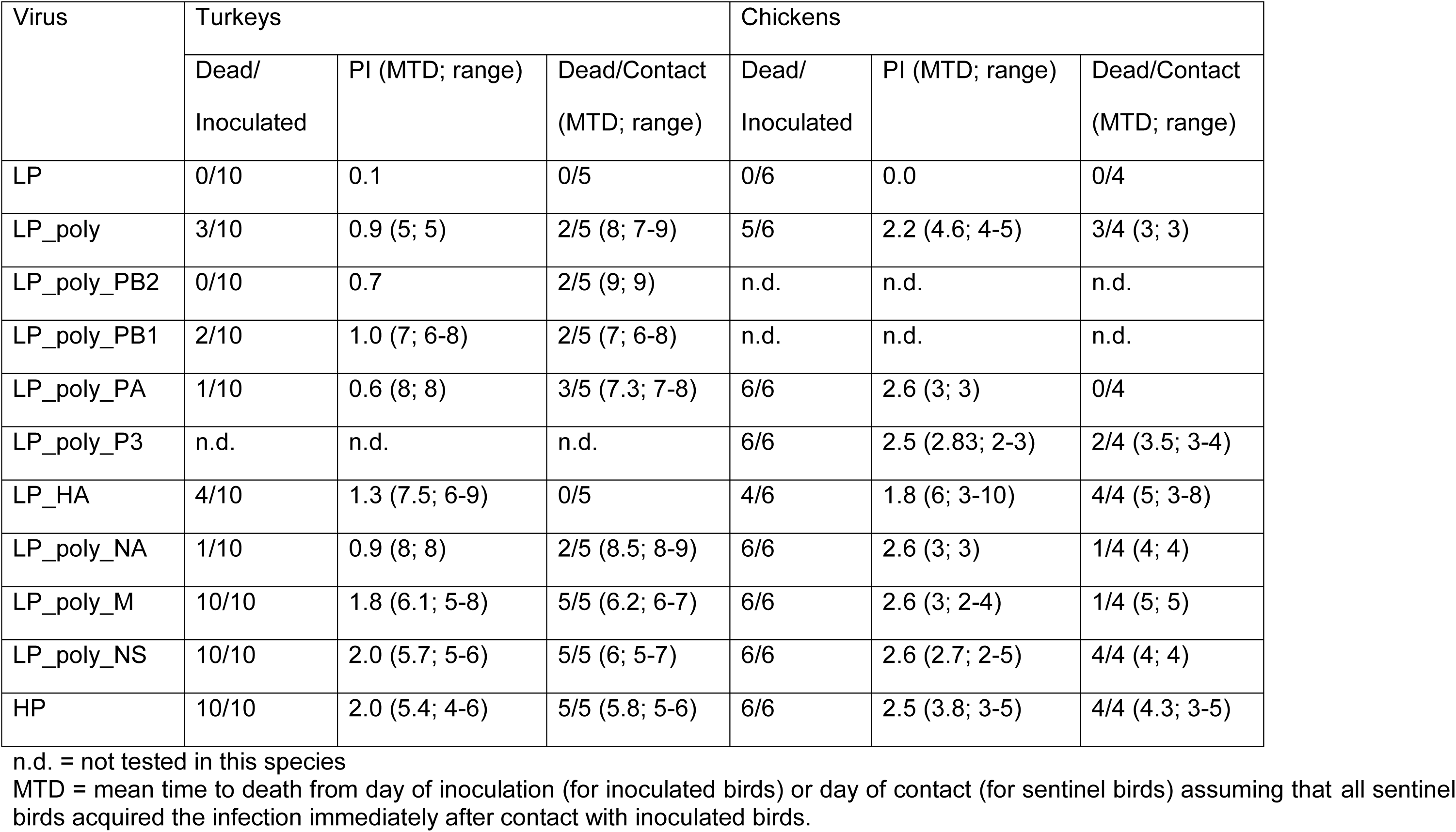
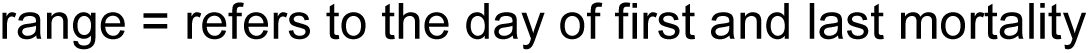
Virulence of recombinant viruses in primarily inoculated or contact chickens and turkeys.

| Virus | Turkeys |  |  | Chickens |  |  |
| --- | --- | --- | --- | --- | --- | --- |
|  | Dead/<br>Inoculated | PI (MTD; range) | Dead/Contact<br>(MTD; range) | Dead/<br>Inoculated | PI (MTD; range) | Dead/Contact<br>(MTD; range) |
| LP | 0/10 | 0.1 | 0/5 | 0/6 | 0.0 | 0/4 |
| LP_poly | 3/10 | 0.9 (5; 5) | 2/5 (8; 7-9) | 5/6 | 2.2 (4.6; 4-5) | 3/4 (3; 3) |
| LP_poly_PB2 | 0/10 | 0.7 | 2/5 (9; 9) | n.d. | n.d. | n.d. |
| LP_poly_PB1 | 2/10 | 1.0 (7; 6-8) | 2/5 (7; 6-8) | n.d. | n.d. | n.d. |
| LP_poly_PA | 1/10 | 0.6 (8; 8) | 3/5 (7.3; 7-8) | 6/6 | 2.6 (3; 3) | 0/4 |
| LP_poly_P3 | n.d. | n.d. | n.d. | 6/6 | 2.5 (2.83; 2-3) | 2/4 (3.5; 3-4) |
| LP_HA | 4/10 | 1.3 (7.5; 6-9) | 0/5 | 4/6 | 1.8 (6; 3-10) | 4/4 (5; 3-8) |
| LP_poly_NA | 1/10 | 0.9 (8; 8) | 2/5 (8.5; 8-9) | 6/6 | 2.6 (3; 3) | 1/4 (4; 4) |
| LP_poly_M | 10/10 | 1.8 (6.1; 5-8) | 5/5 (6.2; 6-7) | 6/6 | 2.6 (3; 2-4) | 1/4 (5; 5) |
| LP_poly_NS | 10/10 | 2.0 (5.7; 5-6) | 5/5 (6; 5-7) | 6/6 | 2.6 (2.7; 2-5) | 4/4 (4; 4) |
| HP | 10/10 | 2.0 (5.4; 4-6) | 5/5 (5.8; 5-6) | 6/6 | 2.5 (3.8; 3-5) | 4/4 (4.3; 3-5) |

n.d. = not tested in this species
MTD = mean time to death from day of inoculation (for inoculated birds) or day of contact (for sentinel birds) assuming that all sentinel birds acquired the infection immediately after contact with inoculated birds.
range = refers to the day of first and last mortality

### In turkeys, the M and NS segments increased virus dissemination across various tissues and exacerbated histopathological lesions without inducing endotheliotropism

To assess virus distribution and microscopic lesions in inoculated turkeys, we performed Immunohistochemistry (IHC) and histological examination. IHC revealed AIV antigen in the parenchyma of the pancreas, brain, kidney, cecal tonsils, and spleen, but not in the liver, duodenum, or skin. The highest antigen expressions were detected in the pancreas and brain, with HP showing the highest levels, followed by LP_poly_M and LP_poly_NS (Figure 1G, Figure 2). Only HP, LP_poly, LP_poly_NS, and LP_poly_M were found in the heart and thymus, while antigen presence in the lung was restricted to HP. Additionally, only HP, LP_poly_M, and LP_poly_NS induced lymphatic necrosis, apoptosis, and/or depletion in the cecal tonsils (Figure 1H). LP_poly and HP caused necrosis in the bursa of Fabricius, while LP_poly_M and LP_poly_NS, similar to HP, caused necrosis and lymphatic depletion in the thymus and spleen. The severity of necrotizing inflammation correlated with antigen concentration (Figure 1I). To determine whether the absence of endotheliotropism in H7N7-infected turkeys was influenced by the inoculation route, turkeys were inoculated intravenously (IV) with LP or HP viruses, followed by necropsy at 4 dpi. IHC analysis confirmed the absence of endotheliotropism in all organs. Overall, the M and NS segments increased virus replication and lesion severity in various organs caused by LP_poly, but endotheliotropism was not observed for any of the investigated viruses.

**Figure 2:**
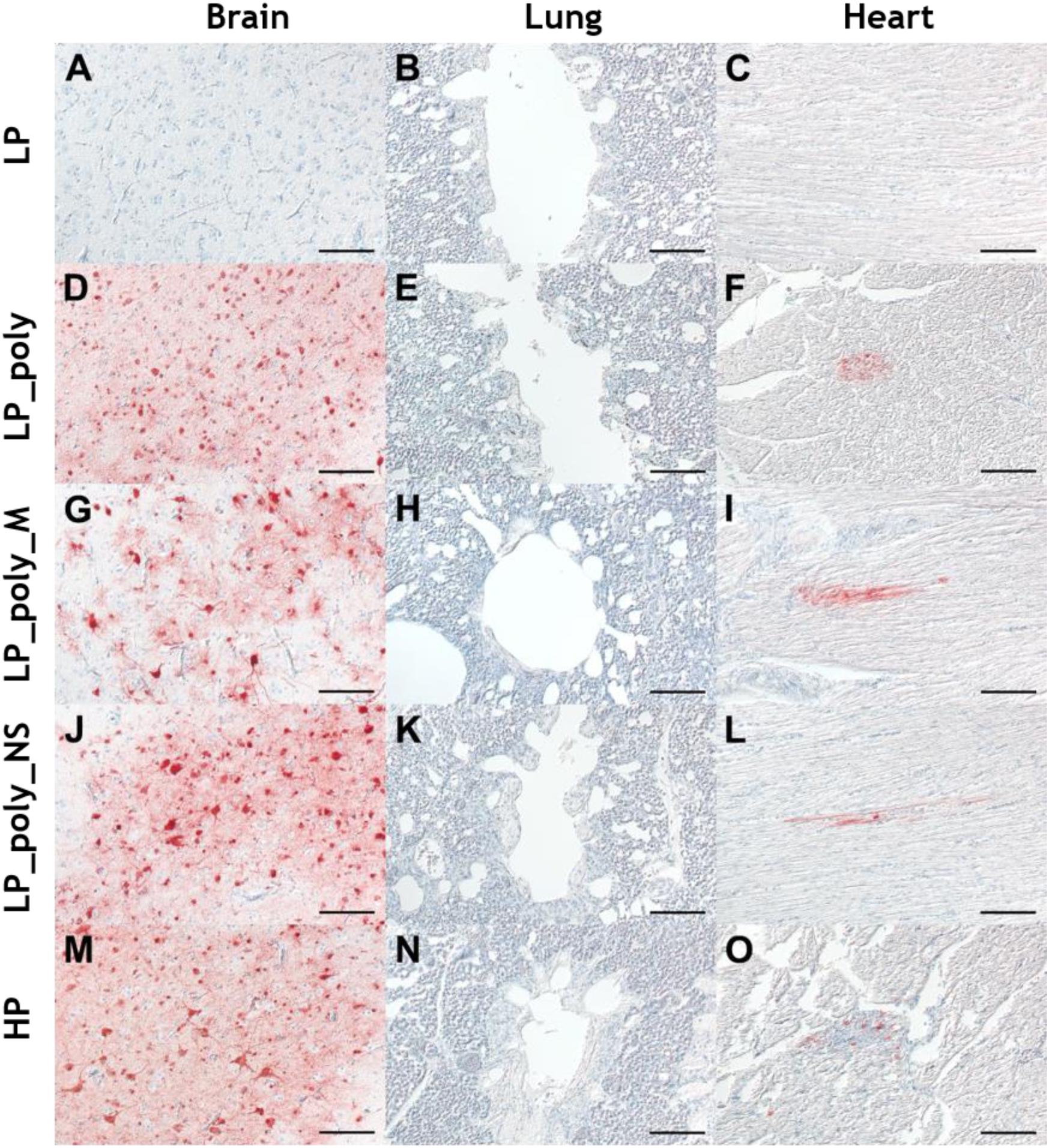
Detection of Influenza A matrix-1 protein (M1) in brain, lung and heart of selected viruses in turkeys. Distribution of influenza A virus matrix-1 protein in different organs in turkeys at 4 days post infection with LP (A-C), LP_poly (D-F), LP_poly_M (G-I), LP_poly_NS (J-L), and HP (M-O). Immunohistochemistry; avidin-biotin-peroxidase complex method; 3-amino-9-ethyl-carbazol (red-brown); hematoxylin counterstain (blue); Nomarski contrast; Bars = 50 µm.

### In chickens, pCS and NS segment contribute to virulence

We inoculated chickens oculonasally and added naïve chickens 1-day post-inoculation to each group to assess direct chicken-to-chicken contact transmission. All HP-inoculated chickens succumbed to infection within five days, with an MTD of 3.8 days and PI of 2.5 (Table 2, Figure 3A). Inoculation with LP_poly resulted in the death of 5/6 chickens, with a PI of 2.2 and MTD of 4.6 days. The surviving bird exhibited moderate clinical signs (e.g., depression, ruffled feathers, diarrhoea, without neurological symptoms), survived until the end of the experiment. Reassorting LP_poly with other HP gene segments (excluding the complete HA) resulted in the death of all inoculated chickens, with comparable PI values of 2.6 and MTD ranging from 2.7 to 3.0 days (Table 2). All co-housed chickens in the HP, LP_poly_NS, and LP_HA groups died, while few or no contact chickens in other groups died (Table 2, Figure 3B). Chickens inoculated with LP_poly or HP showed significantly higher viral RNA excretion via the oral route (p < 0.032) compared to LP inoculated animals. LP_poly, LP_poly_NS and HP were excreted at the highest levels in the OP and CL swabs of contact chickens (Figure 3 E,F). In contrast to the surviving inoculated chickens with LP_poly and LP_HA, no seroconversion occurred in the surviving contact chickens by the end of the experiment, indicating poor virus transmission in the other groups. Together, these findings highlight the pCS as the primary virulence determinant in inoculated chickens, with the NS segment playing an additional role in virus virulence.

**Figure 3:**
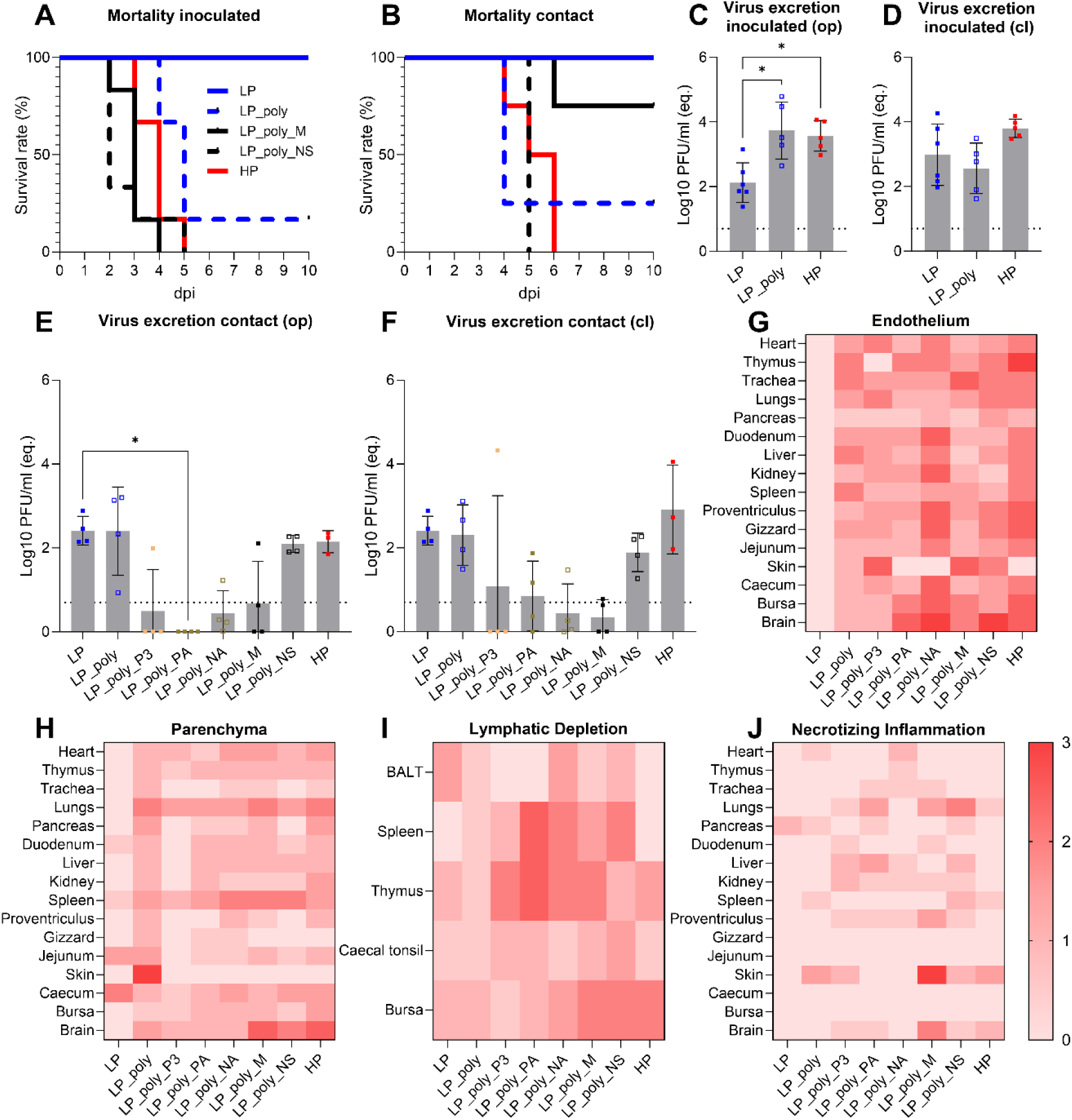
Clinical examination, virus excretion, tissue distribution and microscopic lesion in chickens. Shown are the mortality of inoculated and contact chickens (A, B), virus RNA levels expressed as equivalent log10 plaque forming units/ml in OP and CL swabs at 4 dpi from inoculated (C, D) and contact chickens (E, F). Bars indicate the arithmetic mean of log-transformed data, with individual values represented as dots and standard deviations shown as error bars. The RT-qPCR detection limit is marked by dashed lines. Statistical differences in virus excretion compared to LP-inoculated birds were calculated with Kruskal-Wallis test and p <0.05 was considered significant (C-F). Shown is the median score for distribution of M1 antigen in inoculated chickens at 4 dpi in endothelial cells (G), the parenchyma (H), necrosis in lymphatic organs and tissues (I) and severity of necrotizing inflammation (J).

### In chickens, pCS and NA segment were critical for endothelial infection, while NS and M segments contribute to virus replication and necrosis in the brain

In stark contrast to turkeys, AIV antigen from all viruses with pCS was detectable in the endothelial cells of all organs examined (Figure 3G). Notably, HP and LP_poly_NA showed the highest antigen expression in endothelial cells, with LP_poly_NA exhibiting the widest distribution of the brain. Reassortants with M, NS, and PA segments also facilitated widespread virus distribution, although at lower levels compared to HP and LP_poly_NA. In the parenchyma, HP antigen was present in all examined organs, with the highest expression observed in the brain, lungs, spleen, kidneys, heart, and cecum (Figure 3H). In the brain, LP_poly_M and LP_poly_NS exhibited antigen levels comparable to HP, while other reassortants showed significantly lower amounts (Figures 3H and 4). Lymphatic depletion in the bursa was the highest for HP, LP_poly_M, and LP_poly_NS. While all viruses except LP_poly_NA, LP_Poly and LP caused brain necrosis, LP_poly_NS also induced necrosis in the liver and pancreas (Figure 3I,J). In summary, the pCS and NA segment were major determinants of endothelial tropism, whereas the NS and M segments significantly contributed to virus replication and necrosis in critical organs such as the brain.

**Figure 4:**
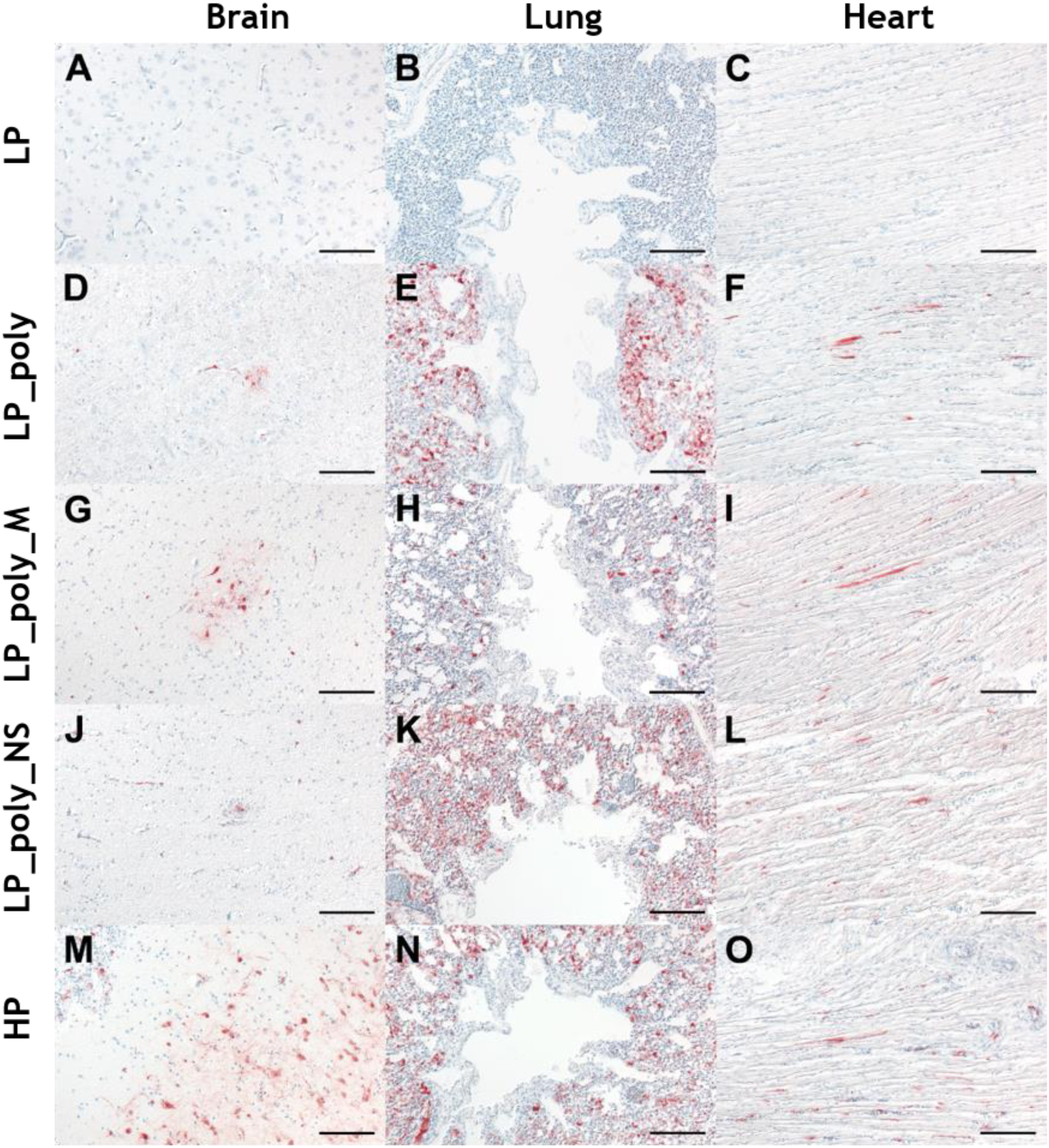
Detection of Influenza A M1 antigen in brain, lung and heart of selected viruses in chickens. Distr ibution of influenza A virus matrix-1 protein in different organs in chickens at 4 days post infection with LP (A-C), LP_poly (D-F), LP_poly_M (G-I), LP_poly_NS (J-L), and HP (M-O). Immunohistochemistry; avidin-biotin-peroxidase complex method; 3-amino-9-ethyl-carbazol (red-brown); hematoxylin counterstain (blue); Nomarski contrast; Bars = 50 µm.

### NS and M segments enhance virus replication with greater impact in turkey cells compared to chicken cells

Given the critical role of the NS and M segments in virulence and replication in chickens and turkeys, we investigated their impact on virus replication in primary turkey embryo kidney (TEK) and chicken embryo kidney (CEK) cells at 1, 8, 24, 48, and 72 hour post-infection (hpi) (Figure 5A,B). In TEK cells, LP virus replicated at the lowest levels across all time points. The insertion of pCS alone significantly enhanced replication only at 8 hpi. However, reassortment with the M or NS segments from the HP virus further boosted replication to levels comparable to the HP virus at all time points. In CEK cells, the addition of pCS, with or without NS and M segments, significantly increased replication levels at 8 hpi. Although reassortment with HP NS or M segments enhanced replication of the LP virus, the levels remained significantly lower than those of the HP virus by ten-times, particularly 48 and 72 hpi. We assessed the cell-to-cell spread of selected viruses. Interestingly, the HP followed by LP_poly_NS and LP_poly_M produced the largest plaques compared to LP and LP_poly (Figure 5C). These findings suggest that the NS and M segments have a variable yet positive impact on replication and spread, with a particularly pronounced effect in turkey cells compared to chicken cells.

**Figure 5:**
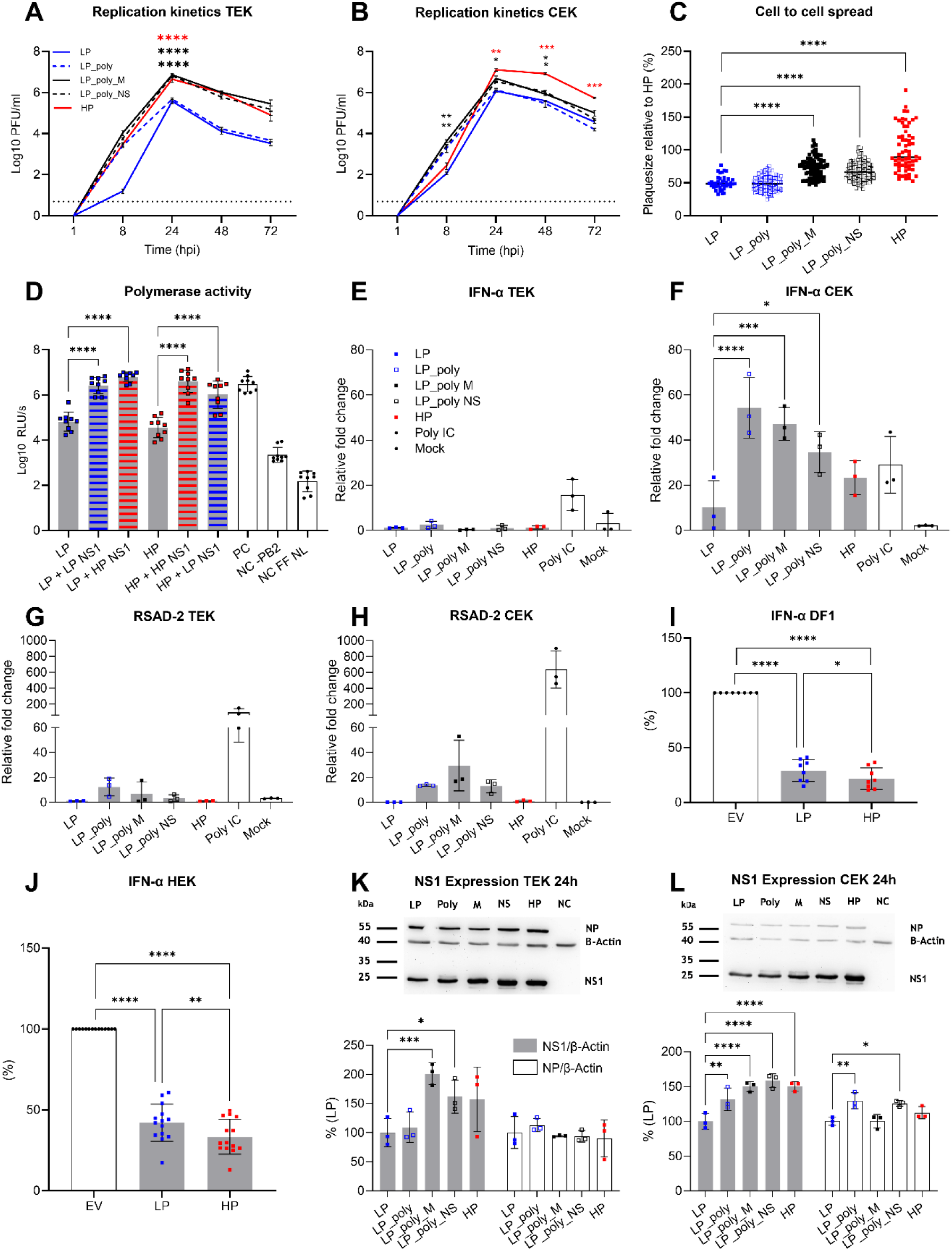
Variation in virus replication, spread, and inhibition of innate immune response. Shown are the results of replication of indicated viruses in TEK cells (A) or CEK (B) expressed as plaque forming units/ml, and cell-to-cell spread in MDCKII expressed as relative plaque size compared to the LP (C). Polymerase activity was determined in a minigenome assay performed on HEK293T cells expressed as log10 Relative Light Units signals. pCAGGS plasmids coding for PB2, PB1, PA and NP from A/swan/Germany/R65/2006 (H5N1) were used as positive controls (PC). As negative control cells were transfected with H5N1 plasmids except PB2 (NC-PB2) or using only Nanoluc luciferase (NL) and firefly luciferase (FF) plasmids (NC FF NL) (D). Transcript levels of IFN-α and RSAD2 in infected TEK and CEK cells. Results were obtained by RT-qPCR and expressed as relative fold change compared to LP. All Ct-values were additionally normalized to GAPDH as a housekeeping gene (E-H). The efficiency of NS1 encoded by pCAGGS on IFN-α induction in chicken DF1 cells and human HEK293T cells via the IRF7 pathway relative to the response in cells transfected with empty pCAGGS vector (EV) (I,J). NS1 expression was quantified using Western blot of infected CEK (K) and TEK (L) cells. Viral proteins were normalized to the housekeeping gene β-Actin. NS1 and NP expression levels are shown in % of the LP NS1 expression. Bars indicate the arithmetic mean of log-transformed data, with individual values represented as dots and standard deviations shown as error bars. The dashed lines in panels A and B refer to the detection limit of the plaque assay. One-way ANOVA with Tukeýs multiple comparisons test was used for statistics on replication kinetics, cell-to-cell spread and polymerase assay. Repeated measures ANOVA was performed as statistical test in the minigenome assay on IFN-induction. Statistical significances in RT-qPCR and Western blot data were determined in an ANOVA with Dunnett’s multiple comparisons test. In all implemented tests a p-value of p<0.05 was regarded as significant. Coloured asterisks in panels A and B refer to the significance compared to LP.

### Polymerase activity of LP and HP was comparable, and increased by NS1 expression

To investigate whether variation in replication levels *in vitro* is due to the polymerase activity, we used a minigenome nano-luciferase reporter assay by transfecting cells with PB2, PB1, PA and NP expressing plasmids with nano-luciferase flanked with HA promotor and firefly reporter genes as control. A comparable level of activity was observed for LP and HP polymerase subunits when co-transfected alone (Figure 5D). A study reported that NS1 enhanced viral polymerase activity of a laboratory PR8 H1N1 virus (23), which may explain the positive impact of HP NS on virus replication *in vivo* and *in vitro* in our studies. Therefore, we did the minigenome assay by adding NS1 pCAGGS expressing vectors to the polymerase subunits (Figure 5D). Interestingly, LP polymerase activity increased significantly when co-expressed with LP NS1 and was further enhanced by HP NS1. Similarly, HP polymerase activity was boosted by HP NS1 and to a lesser extent by LP NS1. Together, these findings highlight that NS1, particularly HP NS1, enhances viral polymerase activity, which may explain its role in driving increased replication and virulence.

### Species-specific interferon modulation by HP and LP NS1 in chicken and turkey cells

The NS1 protein is a key antagonist of the host’s innate immune response, particularly the type I interferon (IFN) response (including IFN-α and IFN-β) and IFN stimulated genes (ISGs). To compare HP and LP NS1 activity in primary chicken and turkey cells, we measured IFN-α mRNA levels, known for its strong antiviral activity, by RT-qPCR 24 hpi (Figure 5 E,F). TEK cells infected with either HP or LP virus exhibited similarly low IFN-α levels, which were significantly lower than those in CEK cells. In CEK cells, HP infection induced a more than tenfold higher mean IFN-α response than LP infection, though this difference was not statistically significant. Notably, compared to LP, LP_poly, LP_poly_M, and LP_poly_NS infections in CEK cells resulted in 44, 36, and 24 folds higher mean IFN-α induction, respectively, a pattern not observed in TEK cells. In a previous study, IFN-stimulated gene expression, particularly RSAD2, differed significantly in H7N1-infected chickens and turkeys (20). To explore this further, we measured RSAD2 mRNA levels in infected cells (Figure 5G,H). RSAD2 expression was significantly lower in TEK than in CEK cells, with no significant differences between viruses in either cell type. To assess the direct effectiveness of NS1 in blocking IFN induction, we expressed HP and LP NS1 using a pCAGGS plasmid and conducted luciferase reporter assays to quantify IFN-α levels, as previously described (24). Since turkey cells could not be transfected, we used chicken DF1 and human HEK293T cells for this experiment. Interestingly, the reporter assay showed that HP NS1, per se, was significantly more effective than LP NS1 in suppressing IFN-α induction in both DF1 and HEK293T cells (Figure 5I,J).

These results indicate that chicken and turkey cells exhibit distinct antiviral responses, with chickens mounting a stronger IFN-α and ISG response than turkeys. While HP and LP NS1 differentially modulate IFN-α responses in chicken cells, additional viral factors, including the pCS and M segments, contribute to IFN regulation.

### Increased NS1 protein expression in turkey and chicken cells

To investigate the expression levels of NS1, we quantified NS1 protein levels in TEK and CEK cells infected with various viruses at 24 hpi. Viral protein expression was normalized to the housekeeping gene β-Actin (Figure 5K,L). Interestingly, NS1 expression was significantly higher in virulent HP and LP_poly_NS compared to LP and LP_poly viruses. Notably, LP_poly_M exhibited elevated NS1 levels in both cell types, despite sharing the same NS segment as LP and LP_poly. We did not find a significant difference on the expression of NP in TEK cells between different viruses, and no impact of LP_poly_M on the expression of NP in CEK cells. This finding suggests that the M segment may independently enhance NS1 expression, highlighting a potential interplay between the M and NS segments in modulating NS1 protein levels.

### Striking variation in virus morphology between chicken and turkey cells

The M segment, encoding the M1 and M2 proteins, plays a critical role in virus budding and morphology. While LP and HP viruses share identical M1 proteins, the HP virus harbours a single M2 mutation (V68L). Given the M segment’s role in virulence and replication, we examined the influence on virus morphology in CEK and TEK cells inoculated with various viruses for 8 hours using electron microscopy (Figure 6). In CEK cells, LP, LP_Poly, and LP_M viruses predominantly formed spherical particles, while HP virus particles were predominately filamentous, suggesting that other gene segments influence virus morphology in chicken cells (Figure 6A,B). In TEK cells, similar to CEK, LP and HP viruses produced predominately spherical or filamentous particles, respectively (Figure 6C,F). Interestingly, unlike in CEK cells, the majority of LP_pCS_M particles in TEK cells were filamentous (Figure 6E). These results were consistently reproducible across different preparations. These findings suggest that the M segment, particularly the M2-V68L mutation, may drive the formation of filamentous virus particles in turkey cells, but not in chicken cells, highlighting a host-specific effect of the M segment on viral morphology.

**Figure 6:**
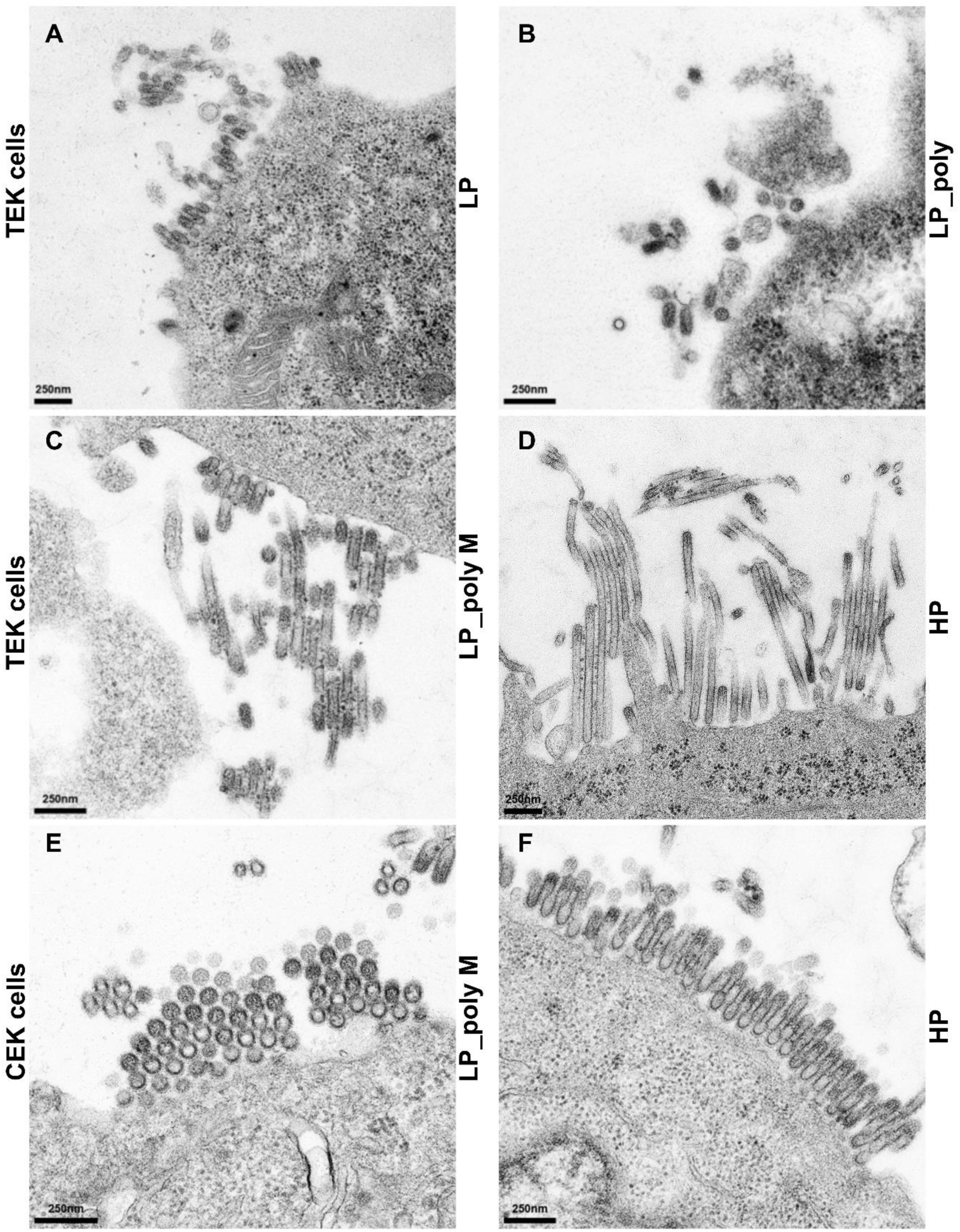
The morphology of virus particles using electron microscopy. Transmission electron microscopy was used to investigate the effect of the pCS and M gene on virus morphology and budding in turkey embryo kidney (TEK) cells (A-D) and chicken embryo kidney (CEK) cells (E+F) infected with indicated viruses for 8 hours at an MOI of 1.

## Discussion

There are significant knowledge gaps regarding viral and host factors driving HPAIV evolution in chickens and turkeys. Virulence determinants in turkeys, the second most industrialized bird species in Western countries, remain largely underexplored. Recent studies showed that a single HA mutation in LPAIV H7N9 (19) or 22-aa deletion in the NA of LPAIV H7N3 (25) enhanced virulence, replication, and transmission in turkeys but not in chickens. Similarly, the incorporation of a pCS alone drove HPAIV H7N1 virulence in turkeys, while additional HA mutations were required in chickens (11, 20). Here, we demonstrate that pCS is the primary virulence determinant of HPAIV H7N7 in chickens but not in turkeys, consistent with our previous findings (22). Notably, HPAIV evolution in turkeys follows two distinct genetic paths – acquiring pCS with HPAIV M or pCS with HPAIV NS –whereas in chickens, only H7N7 with pCS and HPAIV NS was as virulent and transmissible as the HP virus. In simple terms, the evolutionary potential of HPAIV H7N7 from LP ancestor in turkeys would be twice that in chickens. These findings further support a critical role of turkeys in HPAIV evolution and may explain their higher susceptibility to HP-induced morbidity and mortality.

Both host and viral factors shape the distinct pathobiology of HPAIV H7N7 in chickens and turkeys, underscoring key differences in disease progression. In chickens, pCS promoted infection of endothelial cells and systemic dissemination (e.g., heart, thymus, lungs), aligning with its known role in viral spread and endothelial cell replication (26–29). In contrast, pCS alone did not drive endotheliotropism in turkeys, mirroring observations in ducks infected with certain HPAIVs (30, 31). This lack of endothelial tropism in turkeys has also been reported for HPAIV H5N2 (32) and H7N1 (20), and at lower levels in turkeys than in chickens infected with clade 2.3.4.4 H5N8 (33), suggesting fundamental host-virus interactions that limit endothelial involvement. Despite these variations, all viruses were detected in the brain of turkeys, similar to chickens. The potential role of receptor distribution, furin expression, or innate immune responses in endothelial cells remains unexplored. Neuroinvasion via the olfactory nerves (34) may explain the nervous signs in turkeys, contrasting with hematogenous route of infection in chickens (35, 36).

The contribution of NS and M genes to HPAIV pathobiology in turkeys has not been well studied. In chickens, NS1 mutations influenced replication, virulence, or transmission of H5N1 (37) and H5N8 (38). However, NS1 did not significantly affect replication, tissue tropism, or virulence of H7N1 in turkeys, unlike chickens (39, 40). Regardless of endothelial tropism, M and NS segments enhanced virus replication and lesion severity in both species, likely via direct enhancement of viral replication or immune modulation. Finally, NA’s role in endotheliotropism in chickens warrants further study, given its known interaction with host proteases (e.g., plasminogen) and association with neurovirulence in mice (41). NA may facilitate receptor binding by modifying glycans, a mechanism that could differ between chickens and turkeys.

While pCS enhanced single-cycle replication, NS and M segments were essential for sustaining multi-cycle replication in turkey and chicken cells at HP-like levels. Despite no significant difference in polymerase activity between LP and HP, NS1 from both enhanced polymerase function, with HP NS1 having a slightly stronger effect. NS1 suppresses host antiviral responses (42) and modulates polymerase activity by preventing template dissociation, regulating transcription-replication balance, and interacting with NP, ribonucleoproteins, and RNA motifs. It also inhibits host factors like DDX21 and hnRNP-U, stabilizing viral RNA. Additionally, NS1 blocks cellular mRNA processing, leading to nuclear retention of host mRNAs and increasing cap-donor availability for viral transcription (23, 43–45).

NS1 may further enhance replication by directly suppressing the innate immune response. A previous study suggested that some NS1 variants affect AIV replication in turkey cells (46). We found that HP NS1 was significantly more effective than LP NS1 in suppressing IFN-α induction in transfected cells. The HP virus in this study carries the N92D mutation in NS1, while the D92E polymorphism has been linked to interferon response modulation in H5N1-infected chicken and mammalian cells (47, 48). However, little is known about influenza virus interactions with the turkey immune system (46, 49). Our data show distinct antiviral responses in infected chicken and turkey cells, with chickens mounting a stronger IFN-α and ISG response. Previously, we found that HPAIV H7N1-infected turkeys exhibited widespread downregulation of host genes linked to host transcription and translation factors, whereas in chickens, excessive immune activation (immunopathology) may contribute to disease severity and mortality (20).

The M segment encodes M1 and M2 proteins, with M1 playing a crucial role in the virulence and replication of HPAIV H5Nx and H7N9 in chickens (50). M2 is involved at multiple stages of the influenza virus replication cycle, including pH-dependent membrane fusion, blocking autophagy, and facilitating virus budding (51). A V68L mutation in M2 distinguishes HP from LP strains, enhancing virulence, tissue tropism, and excretion, particularly in turkeys, marking the first report of M2’s role in virus fitness in this species. M2 and NS1 cooperatively regulate innate immune responses, including PKR activation, inflammasome signaling, and autophagy, orchestrating programmed cell death to enhance viral replication and release (51–53). This could explain why Lp_poly_M exhibited high IFN and RSAD2 levels alongside robust replication *in vitro* and *in vivo*. However, the impact of the M segment on NS1 expression has not been well investigated. Notably, NS1 has been shown to influence M segment mRNA splicing and nuclear export by interacting with cellular factors (54, 55), thereby promoting viral replication (56). Unfortunately, the commercially available anti-M2 antibody failed to detect the H7N7 M2 protein, preventing assessment of LP and HP NS1 effects on M2 expression. A deeper investigation into the regulation of NS and M splicing and protein expression in LP and HP H7N7 is warranted.

IAV exhibits diverse morphology, ranging from 100 nm spherical particles to filamentous forms exceeding 20 µm (57). Using human isolates or laboratory passaged viruses, morphology was shown to be strain– and cell-specific, influenced by M1/M2 proteins and host factors like the cytoskeleton and Rab11 (58–60). This is the first report to show the variation in virus morphology and size in turkey and chicken cells, driven by the M segment and pCS. Transmission electron microscopy revealed that chicken-derived H2N3 viruses were spherical, whereas duck-derived viruses were elongated despite identical genetics. Filamentous virions are common in human and animal isolates, whereas lab-adapted strains predominantly produce spherical particles, often losing filamentous morphology after serial passage in fowl eggs or MDCK cells (57). Similarly, while filamentous morphology supports IAV spread in guinea pigs, spherical H1N1 variants had a fitness advantage in embryonated eggs (61). The M and NS segments contributed to the spherical H5N1 phenotype with high replication efficiency in human cells and mice (62). Filamentous virions may enhance entry due to a larger HA surface and confer resistance against anti-HA neutralizing antibodies (63). Additionally, M2 interaction with autophagosomal membranes was shown to support filament formation in PR8-H1N1 (64), possibly linking the virus morphology and efficiency to interact with the cellular innate immune response. Further studies are needed to clarify these mechanisms.

This study provides crucial insights into the viral and host factors driving HPAIV evolution in chickens and turkeys. While pCS is the primary virulence determinant in chickens, turkeys follow distinct genetic pathways, with M or NS segments enhancing susceptibility. Host factors such as endothelial tropism and immune responses influence disease progression, with chickens exhibiting stronger endotheliotropism and a more robust innate immune response. Additionally, the interplay between NS1 and polymerase, as well as the contribution of M and NS segments and species-specific virus morphology, further complicates HPAIV evolution. These findings highlight the complex interactions between viral and host factors, supporting a critical role of turkeys in HPAIV evolution.

## Materials and methods

### Viruses and cells

The LPAIV A/chicken/Germany/AR915/2015 (H7N7) (designated hereafter LP) and HPAIV A/chicken/Germany/AR1385/2015 (H7N7) (designated hereafter HP) from the outbreak in Emsland in 2015 were kindly provided by Timm C. Harder. Madin-Darby canine kidney type II cells (MDCKII) were used for virus titration, replication kinetics, assessment of cell-to-cell spread and in combination with HEK 293 T cells for virus rescue. These cell lines were obtained from the Cell Culture Collection in Veterinary Medicine of the FLI. Primary CEK and TEK cells were used for replication kinetics and prepared as previously described (22).

### Generation of recombinant viruses

Cloning of all gene segments of LP and HP viruses into the plasmid pHWS*ccdB* was described previously (22). Rescue of recombinant viruses was done by transfection of targeted plasmids in HEK293T/MDCKII co-cultures using Lipofectamine and OptiMEM (65). Cell supernatant were propagated in specific-pathogen-free (SPF) embryonated chicken eggs (ECE) (Valo BioMedia, Osterholz-Scharmbeck, Germany) according to the standard methods. Nucleotide sequences were generated for all viruses using Sanger sequencing to exclude unplanned mutations. All viruses containing a pCS were handled in biosafety level 3 (BSL3) containments at the FLI.

## Animal experiments

### Ethical statement

All animal experiments in this study were carried out in the BSL3 animal facilities of the FLI following the German Regulations for Animal Welfare after approval by the authorized ethics committee of the State Office of Agriculture, Food Safety, and Fishery in Mecklenburg – Western Pomerania (LALLF M-V).

### Birds

SPF chicken eggs were purchased from Valo BioMedia and commercial turkey eggs were purchased from Juhnke Farm (Tribsees, Germany). Chicks were kept at the FLI animal facilities for hatching. Before infection faecal and swab samples were collected and examined to exclude influenza, parasitic or salmonella infections. Feed and water were added *ad-libitum*.

### Experimental design

At six-week of age birds were allocated into separate rooms at the BSL3 animal facilities at the FLI and kept for two days. At day of infection (d0) six chickens or ten turkeys per group received 10^5^ pfu/ml via oculonasal route in 0.2mL (about 100µL at each side). At 1 dpi four chickens or five turkeys were added to each group to assess virus transmission. The animals were observed for ten days and clinical scoring on a scale from 0 to 3 was done as previously described (11). Briefly, healthy birds were given score (0), sick birds showing one clinical sign (e.g. ruffled feather, diarrhoea, respiratory disorders, nervous manifestation) were given score (1), severely sick birds showed more than one clinical sign were given score (2) and dead birds were given score (3). Severely sick birds were euthanized and scored (3) on the next observation day. The arithmetic mean of clinical signs was calculated for all chickens and the pathogenicity index (PI) was calculated as the sum of the daily arithmetic mean values divided by 10 (the number of observation days) (11). The PI for each virus ranged from 0 (avirulent) to 3 (highly virulent). The scoring was done blindly. On 4 dpi OP and CL swab samples were collected. Viral excretion was determined using generic real-time reverse-transcription polymerase chain reaction (RT-qPCR) targeting the matrix gene after automatic extraction of viral RNA using NucleoSpin**®** 8/96 PCR Clean-up Core Kit (Macherey & Nagel GmbH, Germany) according to the manufacturer instructions. Standard curves were run in each RT-qPCR using serial dilutions of HP virus. Using AriaMix Sotfware Ct-values were plotted against the concentration on the standard curve and then expressed as equivalent log10 PFU/ml. Serum samples collected from all surviving birds at the end of the experiment were tested against anti-NP antibodies using ELISA (IDvet, France) according to the manufacturer instructions.

### Histopathology and immunohistochemistry

At 4 dpi, necropsy was performed under BSL-3 conditions following standard guidelines, and samples from trachea, lungs, pancreas, heart, liver, spleen, kidneys, proventriculus, cecum, duodenum, bursa of Fabricius, thymus and brain from two or three primarily inoculated chickens and turkeys, respectively were collected. Samples were immediately fixed in 4% neutral buffered formaldehyde for 21 days and then embedded in paraffin wax using routine methods. Sections of 5 μm thickness were stained using haematoxylin and eosin. Serial sections were used for immunohistochemistry using the avidin-biotin-peroxidase complex (ABC) method (Vectastain PK-6100; Vector Laboratories, Newark, CA, USA), either a polyclonal rabbit anti-influenza A FPV/Rostock/34-virus-nucleoprotein antiserum, or a primary monoclonal mouse antibody targeting an epitope of the influenza A M1-matrixprotein (clone M2-1C6-4R3 (ATCC® HB-64™), American Type Culture Collection, Manassas, USA), 3-amino-9-ethylcarbazol (AEC) as chromogen (Agilent Technologies, Santa Clara, CA, USA and Nichirei Biosciences Inc., Tokyo, Japan), and hematoxylin counterstain, following the published procedures(66–68). The severity of necrotizing lesions and/or lymphocytic depletion was scored on an ordinal four-step scale (0 = unchanged, 1 = mild, 2 = moderate, 3 = severe), and the distribution of the viral antigen in parenchymal and endothelial cells was evaluated on a four-step scale (0 = none, 1 = focal, 2 = multifocal, 3 = diffuse). as previously described (66).

### Replication kinetics

Primary CEK and TEK cells were infected at an MOI of 0.001. After one hour of incubation at 37°C/ 5% CO2, cells were washed two times with phosphate buffered saline (PBS) and then covered by MEM containing 0.2% BSA (Sigma) and incubated at 37°C/5% CO2. The cells were harvested at 1, 8, 24, 48 and 72 hpi and stored at –80°C until use. The test was done in duplicates for indicated cell culture and repeated three times. Virus titres were determined by plaque assays and expressed as mean titres and standard deviation of all values.

### Plaque assay

Plaque assay was used for titration of viruses in replication kinetics and to estimate the cell-to-cell spread of different recombinant viruses. MDCKII cells were washed once with PBS. The virus solutions were diluted in ten-fold serial dilutions with MEM containing 0.2% BSA and added to the cells for one hour under 37°C/5% CO2 conditions. After one hour the cells were washed twice with PBS and then covered by MEM containing 0.6% BSA which was mixed to equal parts with BactoTM Agar. For the growth of LPAIV, 2 μg/mL of N-tosyl-L-phenylalanine chloromethyl ketone (TPCK)-treated trypsin (Sigma Aldrich, Germany) was added. The plates were incubated for three days at 37°C and fixed using 10% formaldehyde containing 0.1% crystal violet. Viral titres were expressed as plaque forming units per ml (log10 PFU/ml). For the measurement of the plaque size Nikon NIS-Elements software was used. The mean plaque size for each recombinant virus (about 50 plaques each) was expressed as a percentage of the plaque size to the parent recombinant HP virus.

### Mini-genome assay

To measure the polymerase activity of the LP and HP viruses, PB2, PB1, PA, NP and NS1 genes were cloned in pCAGGS plasmid kindly provided by Stefan Finke, FLI, Germany. Plasmids encoding Nanoluc luciferase (NL) and firefly luciferase (FF) flanked by influenza NS1 non-coding regions were kindly provided by Ahmed Mostafa and Stephan Pleschka, JLU, Gießen, Germany. Briefly, HEK293T in 24-well plates were transfected with 250ng each of indicated pCAGGS plasmid combinations in addition to 25ng nanoluc luciferase and 6,25ng firefly pCAGGS coding plasmids using Lipofectamine 2000. After 24 h, cells were lysed using lysis buffer and frozen overnight for better cell disruption. Luciferase activity was determined by NanoGlo® luciferase assay system (Promega) and PJK Beetle-LongGLOW juice according to the manufacturer’s guidelines and normalized relative to the Firefly luciferase activity. The assay was done in triplicates and repeated three times. Results are expressed as average and standard deviation log10 Relative Light Units (RLUs). pCAGGS plasmids coding for PB2, PB1, PA and NP from A/swan/Germany/R65/2006 (H5N1) were used as a positive control. As negative controls cells were transfected with H5N1 plasmids except PB2 and NL/FF plasmids.

### Interferon and ISGs mRNA RT-qPCR

The impact of NS1 mutations on interferon (IFN) and interferon-stimulated genes was analysed using real-time RT-qPCR specific primers as previously described (42–44). Briefly, TEK and CEK cells were infected with the indicated viruses at an MOI of 0.001. After 24 hours, cells were harvested and centrifuged at 10,000 rpm for 10 minutes. Pellets were washed with PBS and processed for RNA isolation using TRI Reagent (Sigma-Aldrich) mixed with PBS in a 1:3 ratio. Following vigorous mixing and a brief spin, 200 µl of chloroform (Carl Roth, Karlsruhe, Germany) was added, and the mixture was incubated at room temperature for 10 minutes before centrifugation at 13,000 rpm for 10 minutes at 4°C. The aqueous phase was transferred to a fresh tube with 600 µl of ≥99.8% ethanol (Carl Roth, Karlsruhe, Germany), mixed, and centrifuged. The lysate was applied to a RNeasy Mini spin column (QIAGEN) and processed per the manufacturer’s protocol. RNA concentration was measured using a Nanodrop photometer, and all samples were adjusted to 10 ng/µl. The following targets were amplified and quantified via RT-qPCR: IFN-α, and RSAD2. Chicken IFN-α was detected using the SuperScript™ III One-Step RT-PCR System with Platinum™ Taq (Thermo Fisher Scientific). Turkey IFN-α, Mx1, and RSAD2 were detected with the SensiFAST™ SYBR® No-ROX One-Step Kit (Meridian Life Science). Transcript levels were normalized to GAPDH and expressed as fold changes relative to the mock control.

### Western Blot

CEK and TEK cells were infected with an MOI of 0.1. Cells were harvested after 24 hours at 37°C. After washing with PBS and centrifugation at 10000rpm/10 minutes the cell pellets were suspended in Laemmli buffer (Serva) and PBS (1:1) and incubated at 95°C for 10 minutes. The cells were centrifuged at 10000rpm for 5 minutes. The proteins were separated from cell lysate by sodium dodecyl sulphate 12% polyacrylamide gels and then electrotransferred to nitrocellulose membranes using a TransBlot cell (BioRad). A monoclonal antibody, derived from the FLI biobank against NS1 or polyclonal rabbit serum against NP were used to detect the NS1 or NP, respectively. Beta-actin (Clone AC74, Sigma-Aldrich) was used as housekeeping protein. Peroxidase conjugated chicken-specific rabbit IGY^++^ or anti-mice IgG (Dianova, Hamburg, Germany) at a concentration of 1:20000 in TBS-T. Immunodetection was achieved by chemiluminescence using Supersignal West Pico chemiluminescent substrate kit (Pierce, ThermoScientific, Rockford, IL, USA) and images were captured using a Bio-Rad VersaDoc Imaging System and Quantity One software. Image J Software was used to analyse the concentration of the NS1 protein compared to the NP and normalized to beta-actin.

### Morphology and budding

The impact of M gene on virus morphology was studied using electron microscopy after infection of CEK and TEK as previously done (69). Briefly, confluent cells in T75 flasks were infected with LP, LP-poly, HP or LP-poly_M for 6 or 8 hours at MOI of 1. Cells were fixed for 30 minutes using 0.5% glutaraldehyde buffered in 0.1 M sodium cacodylate (pH 7.2) (Merck, Germany). Harvested cells were pelleted by low-speed centrifugation and embedded in low-melting-point agarose (Biozym, Germany). Gel slices were then fixed in 1% aqueous OsO4 (Polysciences Europe, Germany). Fixed slices were stained using uranyl acetate followed by stepwise dehydration in ethanol and cleared in propylene oxide, embedded in Glycid Ether 100 (Serva, Germany), and polymerized at 59°C for 4 days. Blocking of samples was done using 0.5 M NH4Cl in PBS for 60 min. Samples were washed with PBS and stained with 0.5% aqueous uranyl acetate for 15 min. Dehydration in ethanol was done under progressive lowering of temperature. Samples were then embedded in the acrylic resin Lowicryl K4M (Lowi, Germany) at −35°C and polymerized by UV light. Ultrathin sections were labelled after blocking of surfaces with 1% cold-water fish gelatin–0.02 M glycine–1% bovine serum albumin fraction V (Sigma, Germany) in PBS by either overnight incubation at 4°C or 2 h of incubation at room temperature with monoclonal anti-NP antibodies (clone HB-65) diluted in PBS-BSA and excess antibodies were removed by washing. Ultrathin sections were examined with a model 400T electron microscope (Philips, The Netherlands) (69).

### Sequence analysis

Analysis of sequences in this study was done using Geneious® software suite v. 8.1.3 (Biomatters, Auckland, New Zealand) to compare mutations in the HP compared to the LP and aligned using MAFFT (70).

### Statistics

An ANOVA with post hoc Tukey tests were utilized to compare replication kinetics, cell-to-cell spread and polymerase activity in the minigenome assay. The statistical test used in the IFN-induction assay was Repeated measures ANOVA. Dunnett’s multiple comparisons test was performed for statistics on expression data of viral proteins in Western blot and RNA-levels of IFN-α and RSAD2. Statistical differences between and within groups for amount of virus excretion and plaque size were evaluated using Kruskal-Wallis test and One-way ANOVA with Tukeýs multiple comparisons test. A p-value of p < 0.05 was considered to be significant; and all analysis was done using GraphPad Prism version 8.1.0.325 (71).

## Acknowledgement

This work was partially supported by grants from the Deutsche Forschungsgemeinschaft (DFG; AB567 and DFG VE780/1-1), DELTA-FLU, Project ID: 727922, an ERA-NET Grant Agreement n° 862605 (ICRAD Flu-Switch) to E.M. Abdelwhab. The work was also funded by the European Union under grant agreement (101084171) – (Kappa-Flu). Views and opinions expressed are however those of the author(s) only and do not necessarily reflect those of the European Union or REA. Neither the European Union nor the granting authority can be held responsible for them. The funders had no role in study design, data collection and analysis, decision to publish, or preparation of the manuscript. The authors are grateful to Nadine Bock, and Laura Timm for laboratory technical assistance, to Bärbel Hammerschmidt, Frank Klipp, Mathias Jan, Doreen Fielder, Bärbel Berger, Thomas Moertiz, and Ralf Henkel for their support in the animal experiments and to Silvia Schuparis for histological preparations. Timm C. Harder is thanked for providing the viruses, Stephan Pleschka, Ahmed Mostafa and Stefan Finke are thanked for providing different pCAGGS plasmids. Sven Reiche and Daniel Marc are thanked for providing the monoclonal antibodies against M1 and NS1, respectively.

